# Modeling spatio-temporal annual changes in the probability of human tick-borne encephalitis (TBE) occurrence across Europe

**DOI:** 10.1101/2024.10.18.619031

**Authors:** Francesca Dagostin, Diana Erazo, Giovanni Marini, Daniele Da Re, Valentina Tagliapietra, Maria Avdicova, Tatjana Avšič –Županc, Timothée Dub, Nahuel Fiorito, Nataša Knap, Céline M. Gossner, Jana Kerlik, Henna Mäkelä, Mateusz Markowicz, Roya Olyazadeh, Lukas Richter, William Wint, Maria Grazia Zuccali, Milda Žygutienė, Simon Dellicour, Annapaola Rizzoli

## Abstract

**Introduction:** Tick-borne encephalitis (TBE), caused by tick-borne encephalitis virus (TBEV), is a zoonotic disease that can cause severe neurological symptoms. Given the increasing number of reported human TBE cases in Europe, a spatio-temporal predictive model to infer the year-to-year probability of human TBE occurrence across Europe at the regional and municipal administrative levels was developed.

**Methods:** The distribution of human TBE cases at the regional (NUTS-3) level during the period 2017-2022, was derived by using data provided by the European surveillance system (TESSy, ECDC), and at the municipal level by using data from Austria, Finland, Italy, Lithuania, and Slovakia. The probability of presence of human TBE cases at the regional and municipal levels for the period 2017-2024 was modeled with a boosted regression trees model, including covariates that affect both the natural hazard of virus circulation and human exposure to tick bites.

**Results:** Areas with the highest probability of human TBE infections are primarily located in central-eastern Europe, the Baltic states, and along the coastline of Nordic countries up to the Bothnian Bay. Our results also highlight a statistically significant rising trend in the probability of human TBE infections not only in north-western, but also in south-western European countries, offering a spatio-temporal predictive framework for the assessment of areas where human TBE infection are most likely to occur. The model showed good predictive performance, with a mean AUC of 0.85 (SD = 0.02), sensitivity of 0.82, and specificity of 0.80 at the regional level, and a mean AUC of 0.82 (SD = 0.03), sensitivity of 0.80, and specificity of 0.69 at the municipal level.

**Discussion:** With ongoing climate and land use changes, the number of human TBE cases is likely to increase and expand into new areas, as trends are already indicating. This underscores the need for predictive models that can help prioritize intervention efforts. The approach adopted, by leveraging lagged covaries, enables timely one-year-ahead predictions, thus supporting surveillance, prevention, and control of human TBE infections by public health authorities.

**Statements:** *Ethical statement:* Ethical approval was not needed.

*Funding statement:* This project has received funding from the European Union’s Horizon 2020 research and innovation programme under grant agreement No 874850 and is catalogued as MOOD 081. The contents of this publication are the sole responsibility of the authors and don’t necessarily reflect the views of the European Commission.

*Conflict of interest:* None.

*Authors’ contributions:* Francesca Dagostin: Conceptualization, Methodology, Data Curation, Formal Analysis, Writing - Original Draft. Diana Erazo: Conceptualization, Methodology, Writing - Review & Editing. Giovanni Marini: Conceptualization, Methodology, Writing - Review & Editing. Daniele Da Re: Conceptualization, Methodology, Writing - Review & Editing. Valentina Tagliapietra: Conceptualization, Methodology, Writing - Review & Editing. Maria Avdicova: Resources, Writing - Review & Editing. Tatjana Avšič – Županc: Resources, Writing - Review & Editing. Timothée Dub: Conceptualization, Resources, Writing - Review & Editing. Nahuel Fiorito: Resources, Writing - Review & Editing. Nataša Knap: Resources, Writing - Review & Editing. Céline M. Gossner: Resources, Writing - Review & Editing. Jana Kerlik: Resources, Writing - Review & Editing. Henna Mäkelä: Resources, Writing - Review & Editing. Mateusz Markowicz: Resources,Writing - Review & Editing. Roya Olyazadeh: Resources, Writing - Review & Editing. Lukas Richter: Resources, Writing - Review & Editing. William Wint: Resources, Writing - Review & Editing. Maria Grazia Zuccali: Resources, Writing - Review & Editing. Milda Žygutiene: Resources, Writing - Review & Editing. Simon Dellicour: Methodology, Writing - Review & Editing. Annapaola Rizzoli: Conceptualization, Methodology, Writing - Review & Editing.

*Data availability:* The data that support the findings of this study were provided by ECDC, Azienda Provinciale per i Servizi Sanitari Provincia Autonoma di Trento (APSS), Unità Locale Socio Sanitaria Dolomiti (ULSS N.1 Dolomiti), Public Health Authority of the Slovak Republic, Austrian Agency for Health and Food Safety (AGES), Finnish Institute for Health and Welfare (THL), National Public Health Center under the Ministry of Health (Lithuania) and University of Ljubljana. Restrictions apply to the availability of these data, which were used under license for the current study, and so are not publicly available. The interactive risk maps can be explored in detail at https://mood-platform.avia-gis.com.

*Disclaimer:* The views and opinions of the authors expressed herein do not necessarily state or reflect those of ECDC. The accuracy of the authors’ statistical analysis and the findings they report are not the responsibility of ECDC. ECDC is not responsible for conclusions or opinions drawn from the data provided. ECDC is not responsible for the correctness of the data and for data management, data merging and data collation after provision of the data. ECDC shall not be held liable for improper or incorrect use of the data.

## Introduction

Tick-borne encephalitis (TBE) is a severe viral infection of the central nervous system caused by the tick-borne encephalitis virus (TBEV), a member of the Flaviviridae family [1,2]. TBEV is primarily transmitted to humans through bites from infected ticks of the Ixodidae family. However, transmission can also occur through non-vectorial routes of infection, such as the consumption of unpasteurised milk and dairy products from infected ruminants [3], accounting for about 1% of all TBE cases reported annually in the EU/EEA. While mild cases may present flu-like symptoms, severe cases can lead to inflammation of the brain or the membranes surrounding the brain and spinal cord, resulting in potentially life-threatening complications, and long-term neurological deficits in survivors [1,2]. The most widespread TBEV subtype circulating in the EU/EEA is the European subtype (TBEV-Eu), while the Siberian subtype (TBEV-Sib) occurs in some northeastern European countries [4]. In TBEV-Eu infections, 80% generally remain asymptomatic. The remainder usually present with biphasic disease, of which 10% show symptoms with severe neurological sequelae and a mortality rate of 1% to 2% [1]. Since 2012, TBE has been a notifiable disease in the EU/EEA, with mandatory reporting in nineteen countries, voluntary reporting in four (Belgium, France, Luxembourg, and the Netherlands) and ‘not specified’ in one country (Croatia). The increase in TBE cases across Europe - from 2412 in 2012 to 3514 in 2022, coupled with the emergence of new foci of infection in previously non-endemic countries [5,6], has highlighted the need for predictive tools capable of identifying areas where human TBE infections are likely to occur.

Surveillance efforts for TBE aim to track the incidence and distribution of TBE infections in the human population, identify high-risk areas, and assess temporal trends in disease prevalence. Within this context, mapping the suitability for the occurrence of human TBE cases at the finest spatial resolution, ideally at the municipal level, is essential for targeted public health interventions, including awareness and vaccination campaigns. Machine learning techniques, which can handle complex non-linear relationships and different types of predictors, have become increasingly valuable in predicting the potential distribution of vectors and infectious diseases [7]. Such approaches have been successfully applied to model the distribution of diseases like dengue [8], chikungunya [9], and West Nile fever [10,11]. However, for tick-borne diseases, European studies have predominantly focused on predicting the distribution of tick vectors [12,13] or ecological suitability for TBEV in tick or animal host populations [14,15], rather than the probability of occurrence of human infections. In the case of modelling studies focused on human TBE cases, recent publications focus only on limited areas or specific countries, such as Finland [16], Germany [17], or Sweden [18]. At the continental scale, maps that retrospectively depict human TBE notification rates are available but usually have a rather coarse spatial resolution, i.e. at the Country level [5].

Therefore, the development of a modeling framework that estimates the probability of human TBE infections at the finest administrative scale based on easily accessible data on environmental, animal, climatic and anthropogenic factors, represents a step forward towards comprehensive TBE risk estimation and mitigation in Europe. In response, this study presents a novel spatio-temporal modelling framework that provides annual predictions of habitat suitability for human TBE infections across Europe, at both the regional and municipal levels, which will be updated annually before the start of tick questing season. In our model, we considered predictors that reflect both the natural hazard of virus circulation and human exposure to tick bites, a dimension generally lacking in previous models. In fact, the geographical distribution of human TBE infections depends on both hazard factors that drive the local circulation of the TBE virus [19,20], and the likelihood of human encounters with infected ticks [21].

Our specific objectives are to (i) generate annual suitability predictions for human TBE infections at regional and municipal administrative levels, which will be updated annually before the tick questing season, (ii) generate suitability maps for human TBE cases based on these predictions, and (iii) assess spatio-temporal trends in the probability of human TBE occurrence. Overall, this predictive framework will aid in identifying areas suitable for targeted TBEV detection surveys and will support public health authorities in planning surveillance and prevention efforts one year in advance.

## Methods

### Collection of epidemiological data

The observations of reported human cases of TBE were obtained from The European Surveillance System (TESSy), a database for the collection, validation, cleaning, analysis, and dissemination of data for public health activities hosted by the European Centre for Disease Prevention and Control (ECDC). The choice to base our model on notified human TBE cases reflects the availability of systematically collected, centralized data for several European countries across multiple years. Epidemiological data included, where available, the most likely place of infection at “nomenclature of territorial units for statistics” (NUTS) level 3 (which corresponds to small regions for specific diagnosis, according to the NUTS classification system [22]). In our study, we used the human TBE confirmed and probable cases reported to TESSy between 2017 and 2022. To improve spatial accuracy, we excluded patients who acquired the infection abroad or whose place of exposure was unknown or not reported at the NUTS-3 administrative level and we only included countries with sufficient reporting, i.e., that provided the location of infection at the NUTS-3 level for at least 75% of cases notified during the selected period. The 13 countries selected according to these criteria were: Czechia, Denmark, Germany, Greece, Finland, France, Hungary, Italy, Lithuania, Poland, Romania, Slovakia, and Sweden. To complement this dataset, we received data from Austria (Austrian Agency for Health and Food Safety, AGES) and Slovenia (University of Ljubljana). We also included Ireland and Spain, two countries in which, despite compulsory notification, no autochthonous cases were reported during the study period, to account for areas with no recorded presence of TBE.

To further improve our dataset, we also obtained, upon formal request, data reported at a finer administrative level, namely the distribution of human TBE cases in Austrian districts between 2017 and 2022, provided by the Austrian Agency for Health and Food Safety (AGES), and data collected between 2017 and 2022 in the municipalities of Finland (provided by the Finnish Institute of Health and Welfare, THL), Slovakia (provided by the Public Health Authority of the Slovak Republic), Lithuania (provided by the National Public Health Centre under the Ministry of Health) and northern Italy (provided by the local public health agencies, Azienda Provinciale per i Servizi Sanitari Provincia Autonoma di Trento and Unità Locale Socio Sanitaria Dolomiti Belluno). Based on these data sources, we compiled two dichotomous datasets representing the presence and absence of human TBE cases between 2017 and 2022, at the regional (NUTS-3) and municipal levels. Locations identified as “presence” for human TBE, were defined as administrative units reporting at least one confirmed human TBE case per year, while absence locations were defined as the remaining units with zero confirmed cases. It’s important to note that “absence” here pertains solely to the lack of reported human TBE cases, not to the absence of TBEV circulation within ticks and hosts. Thus, regions categorized as “absence” may indeed harbor TBEV within their tick and wildlife populations; however, no human cases have yet been detected, possibly due to lower human exposure to tick bites in these areas.

### Hazard and exposure covariates

We included in our study a comprehensive set of covariates that reflect both hazard and exposure dimensions, as risk assessment relies on the interaction between these two components (Figure 1). Specifically, hazard refers to any potential source of harm, while risk is the probability of harm occurring based on physical exposure to the hazard. Concerning tick-borne diseases, hazard can be defined as the presence or density of infected vectors in the environment, which depends on the interplay of complex ecological and environmental conditions that allow TBEV circulation among ticks and competent hosts [23]. Exposure, on the other hand, relates to human interactions with the environment that increase the likelihood of encountering infected ticks and depends on factors such as human outdoor activities and socio-economic variables [18].

**Figure 1.**
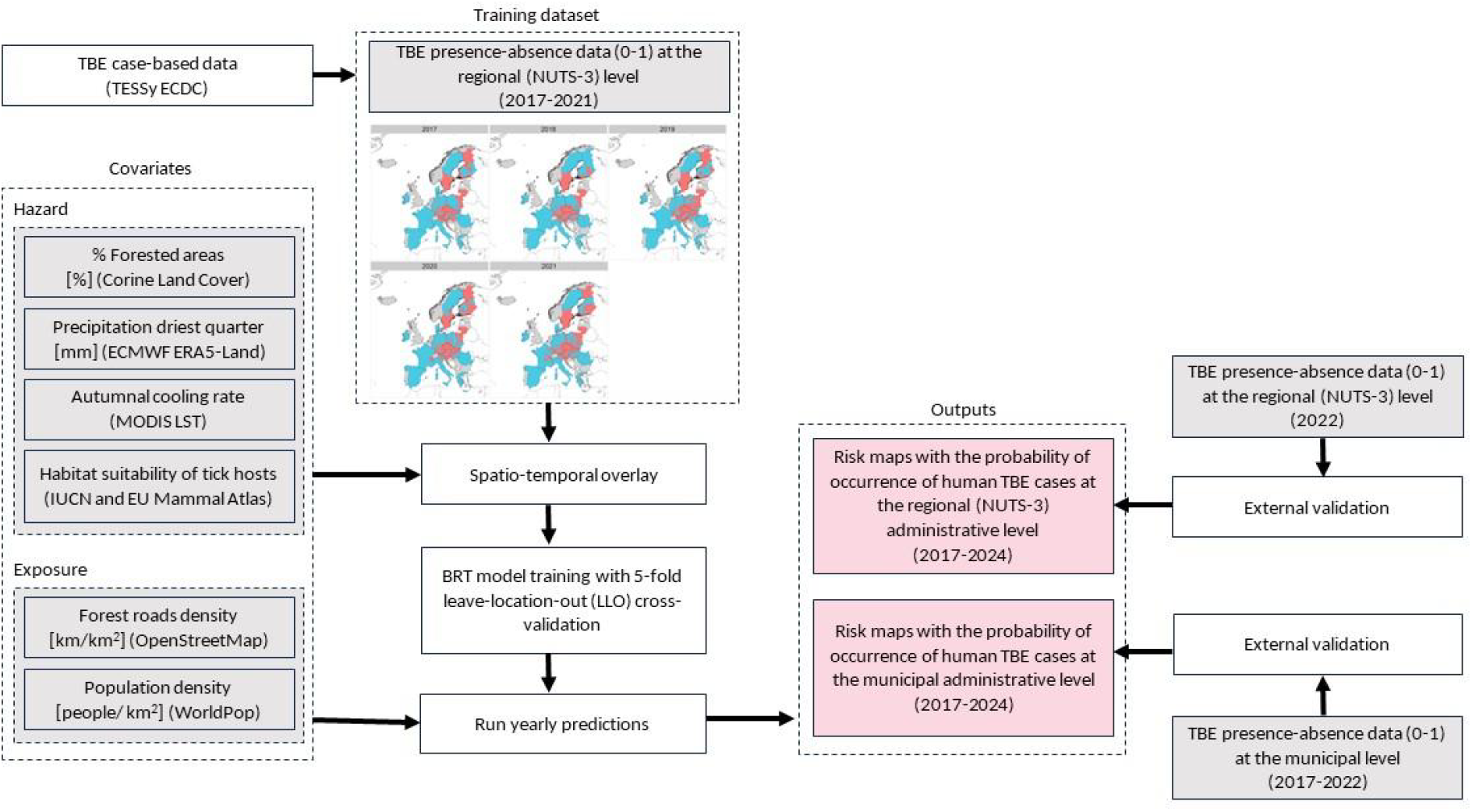
Schematic representation of the modelling framework.

We selected key hazard factors as suggested in [24]. These predictors included: (i) the percentage of forested areas in each area, (ii) the total precipitation of the driest quarter one year prior, (iii) the autumnal cooling rate one year prior, (iv) the weighted probability of presence of selected critical host species (rodents and deer) originally produced using Random Forest and Boosted Regression Trees based spatial modelling techniques [25] (see Supplementary Material for a detailed description of covariates and data sources). The assumption of a constant distribution (presence or absence of key hosts) during the study period was made due to the lack of standardised, high-resolution data on annual and interannual variations in host abundance and density at the continental scale.

To account for human exposure to tick bites, assuming that roads increase access and that forests with roads are more likely to be entered by visitors, we computed the density of forest roads contained by OpenStreetMap as a proxy of accessibility to forested areas [18]. Human population density in each area, derived from the WorldPop dataset [26], was also included. Time-dependent predictors, the autumnal cooling rate and the precipitation of the driest quarter, were computed for each year of analysis with a 1-year lag (2016-2023). All explanatory variables were computed by averaging them for each administrative area included in the dataset, at the regional and municipal administrative levels (see Supplementary Material, Figure S1-S6, for the mapped values of predictors).

### Boosted regression trees modelling framework

We modelled the probability of the presence of human TBE cases using a boosted regression tree (BRT), a machine-learning technique based on regression trees and gradient boosting. BRT offer several advantages over other modelling techniques, including the capacity to handle diverse types of predictor variables, to fit complex nonlinear relationships, and to accommodate missing data [7]. We implemented the BRT algorithm using the “gbm” R package [27] using spatial cross-validation to limit model overfitting. Specifically, we applied a leave-location-out (LLO) resampling method within the ‘mlr3’ R framework [28] in which a test set is selected and all observations that correspond to the same location across multiple time points are omitted from the training sample. To consider spatial autocorrelation, spatial fold selection was defined based on the block generation method described by Valavi and colleagues and implemented in the R package “blockCV” [29]. As BRT builds the trees on random subsamples, we ran 10 independent model replicates and averaged the results.

We trained the BRT model on human TBE presence-absence data reported between 2017 and 2021 at the NUTS-3 spatial level (“regional model”), and assessed the model predictive performance by estimating the area under the receiver operating characteristic curve (AUC), the sensitivity and specificity, and the prevalence-pseudoabsence-calibrated Sørensen’s index (SI_ppc_) [30,31], as the use of the AUC metric alone has been repeatedly criticised in previous works because of its dependence on sample prevalence (i.e., the ratio between the number of presences and number of absences) [32]. As the computation of this index needs binary presence-absence data, whereas our model instead returns predicted probability values, we performed an optimisation procedure by applying a threshold to the model’s predictions to transform them into a presence-absence dataset. The threshold values were varied in the range [0, 1] with a 0.01 step increment, and the threshold that maximised the SI_ppc_ was eventually selected to compute the index (see [33] for the application of a similar approach). Performance metrics were calculated across the ten replicates.

In addition, official TBE records from 2022, reported at the NUTS-3 level, were left out of model training and later used to externally assess the predictive power of the regional model.

Finally, we applied the model trained with human TBE presence-absence data reported between 2017 and 2021 at the NUTS-3 spatial level (as described in the above paragraph) to predictors averaged at the municipal administrative level to obtain a higher resolution output [34]. The reason for applying the regional model at the municipal level was due to the lack of sufficient municipal-level data to develop a model based solely on them. The accuracy of municipal predictions was externally assessed based on the presence/absence dataset compiled at the municipal level for the period 2017-2022 (see Supplementary Material, Figure S7, for the presence/absence dataset of TBE at the municipal level). The model enables the prediction of human TBE suitability on an annual basis, using temporal covariates recorded one year prior, and results will be updated yearly before the start of the tick questing season. For example, the prediction for 2025 could be made by February 2025, based on the covariates recorded in 2024. A schematic representation of the modelling framework is shown in Figure 1.

Finally, linear regressions were performed to identify areas with either an increasing or decreasing statistically significant trend in the probability of human TBE cases over the specified years and across the two administrative scales. Specifically, for each area, we fitted a generalised linear model (GLMs) to the predicted human TBE probability across the years (2017-2024), with the year as the predictor variable (formula: prediction ∼ year). All analyses were performed using R Statistical Software v.4.1.2 [35].

## Results

The current observed spatial distribution of human TBE cases across Europe extends westward to the Ain department of France, southward to northern Italy, eastward to the Balkan countries, and northward to Norway and Finland. Over the period spanning 2017 to 2022, 15 European countries reported at least one confirmed, autochthonous human case with a known place of infection at the regional level. Overall, this dataset comprised a total of 1,658 “presences” in NUTS-3 regions over the period of analysis, spanning 15 European countries (Austria, Czech Republic, Denmark, Germany, Greece, Finland, France, Hungary, Italy, Lithuania, Poland, Romania, Slovakia, Slovenia, and Sweden). The number of NUTS-3 positive regions (*i*.*e*., that notified at least one human TBE case), increased from 255 regions in 2017 to 279 regions in 2022. Within the same timeframe, no human TBE cases were reported from 4,096 other regions within the 15 countries mentioned above, plus Spain and Ireland (Figure 2).

**Figure 2.**
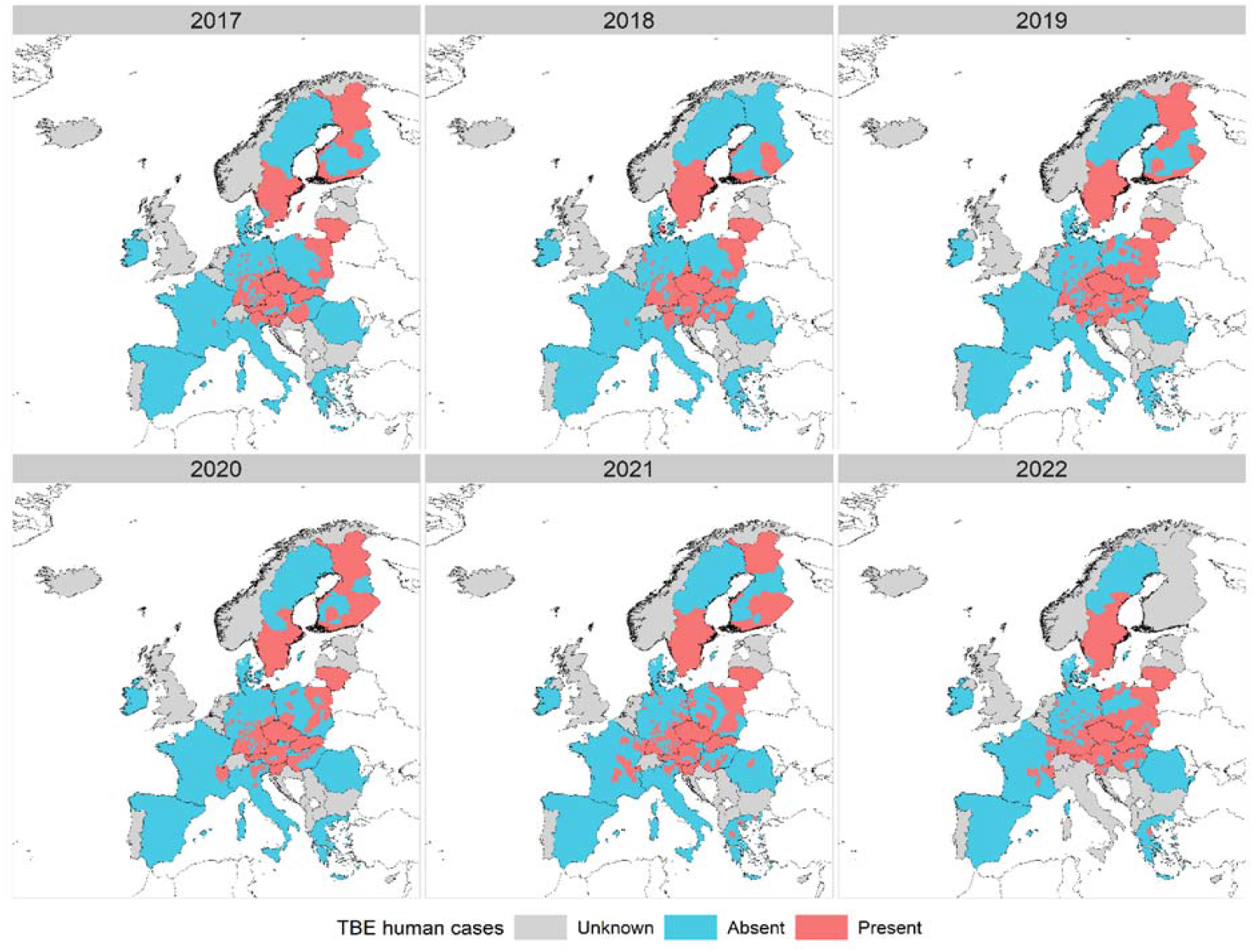
Presence (in red) and absence (in blue) of human TBE cases at the NUTS-3 (regional) administrative level (2017-2022). Areas where human TBE incidence is unknown (either data were not reported, or not reported at the NUTS-3 level) are masked in grey.

### Predictions of the probability of human TBE presence at the regional and municipal level

The predicted probability of occurrence of human TBE cases at the regional level is shown in Figure 3. Figure 4 provides more detailed maps, displaying the same probability at the municipal level. The mean AUC was 0.85 (standard deviation (SD) = 0.02) for the regional model, derived from internal cross-validation, and 0.82 (SD = 0.03) for the model at the municipal scale, derived from external validation. The predictive performance at the regional scale showed good sensitivity (0.82) and specificity (0.80), while specificity decreased when downscaling the results at the municipal scale (0.69). Similarly, the Sorensen’s Index showed a variation from 0.71 (regional model) to 0.56 (municipal model).

**Figure 3.**
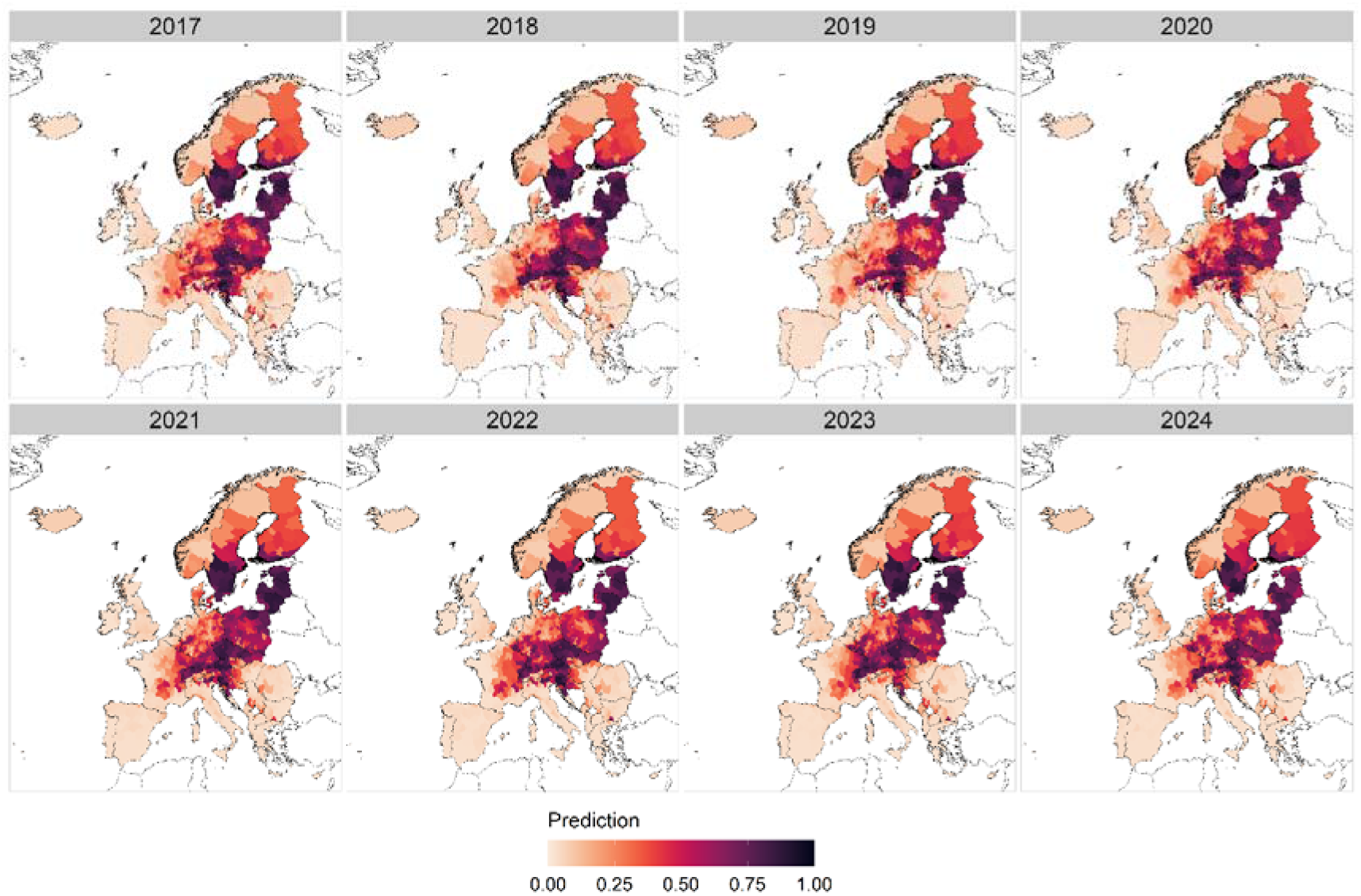
The predicted probability of the occurrence of human TBE cases at the regional (NUTS- 3) level, 2017-2024.

**Figure 4.**
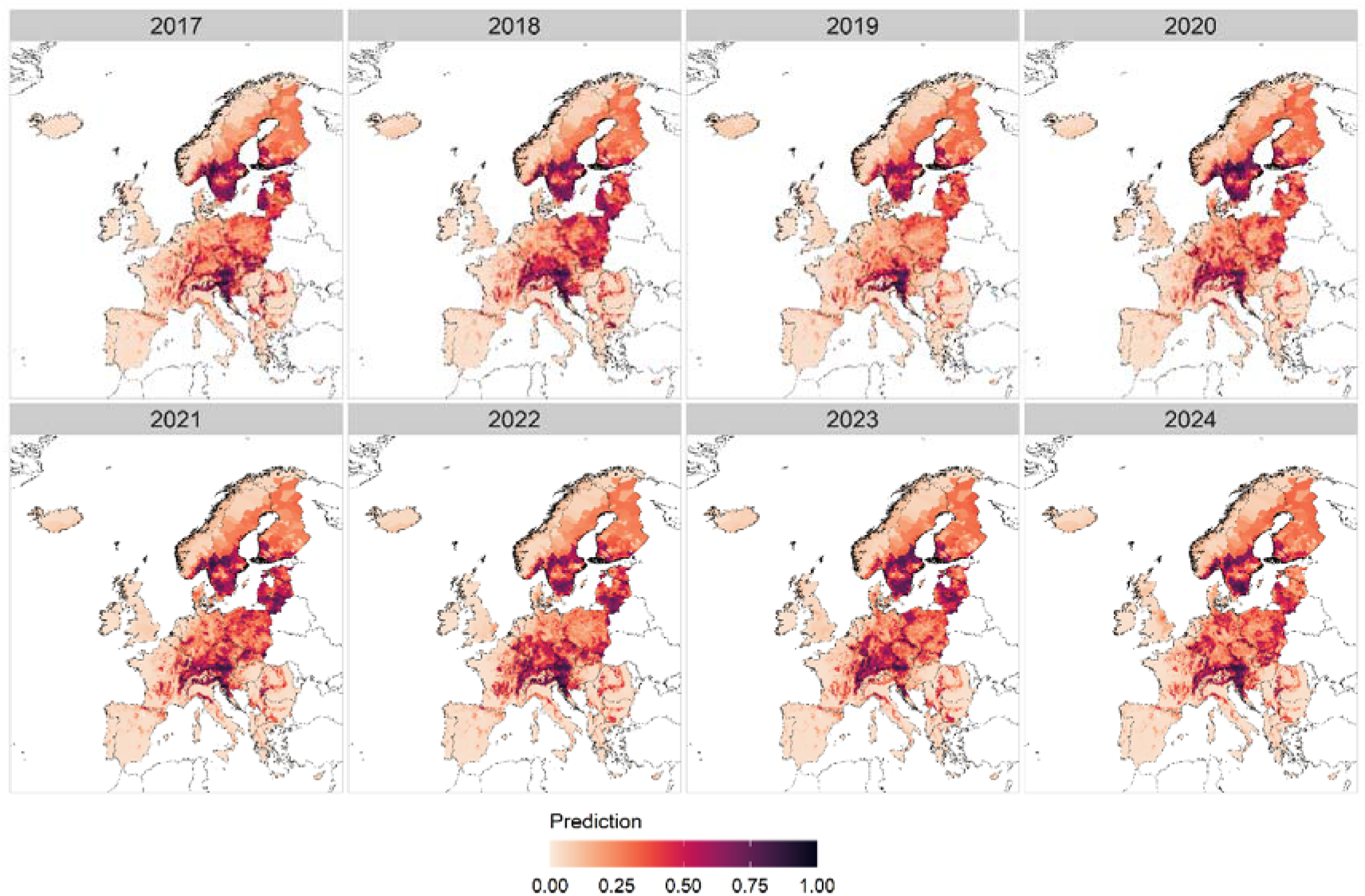
The predicted probability of the occurrence of human TBE cases at the municipal level, 2017-2024.

We also performed external validation on official TBE cases reported during 2022 at the regional level, which were not included in the training dataset. The external validation of regional predictions yielded good results, with an AUC of 0.87, sensitivity of 0.81, specificity of 0.78, and SI_ppc_ of 0.73 (see Supplementary Material, Table S3, for detailed performance metrics).

### Fitted functions and relative importance of variables

BRT results comprise two main elements: fitted functions and the relative importance of variables. Fitted functions can be visualised using partial dependence plots. They show the effect of a predictor on the response after accounting for the average effects of all other variables and were interpreted here as the relative probability of reporting at least one human TBE case at various levels of the predictor variable. The contribution of each environmental factor is given by its relative influence (RI) in the BRT model and is computed as the number of times the variable is selected for splitting a tree, weighted by the squared improvement to the model resulting from each split averaged over all trees [7]. Variables which showed a positive trend are the probability of vertebrate hosts’ presence (RI = 39.44); the density of roads and pathways in forested areas (RI = 21.39); the total precipitation of the driest quarter (RI = 20.19); the proportion of forested areas (RI = 14.85); and population density (RI = 2.37). The autumn cooling rate (RI = 1.77), which is negative by definition and measures the rate of decrease in late summer temperatures, showed a negative trend, meaning that human TBE suitability is higher when such temperature decrease is steeper (see Supplementary Material, Figure S8, for partial dependence plots).

### Short-term variability of predicted probability of human TBE presence in Europe

To examine potential upward or downward trends in the spatio-temporal variation of the predicted probability of occurrence of human TBE cases from 2017 to 2024, we conducted GLM analyses for each specific administrative unit, considering the BRT prediction as the response variable and the year as the predictor variable, and mapped statistically significant coefficients both at the NUTS-3 (Figure 5a) and municipal (Figure 5b) administrative levels. The results suggest a statistically significant increase in TBE probability of presence across central-western European countries (e.g. Switzerland and Germany), with the emergence of new areas suitable for human TBE infections extending in a north-westerly and potentially south-westerly direction (e.g. Belgium, France, and the United Kingdom). Areas with larger significant increases in the predicted probability of human TBE presence can be identified at both regional (Figure 5a) and municipal (Figure 5b) levels and are concentrated in western Germany, southern Belgium, eastern France, and Switzerland (dark red areas), suggesting a non-negligible statistically significant short-term increase in areas where human TBE infections are likely to occur in recent years, and so possibly in the coming years. In contrast, areas of significant decrease in the predicted probability of human TBE cases can be seen in parts of Eastern Europe (purple areas). It is important to note that this decrease only implies a reduction in the predicted probability over time. Human TBE infections may still occur in these areas, but there has been a decreasing trend in recent years.

**Figure 5.**
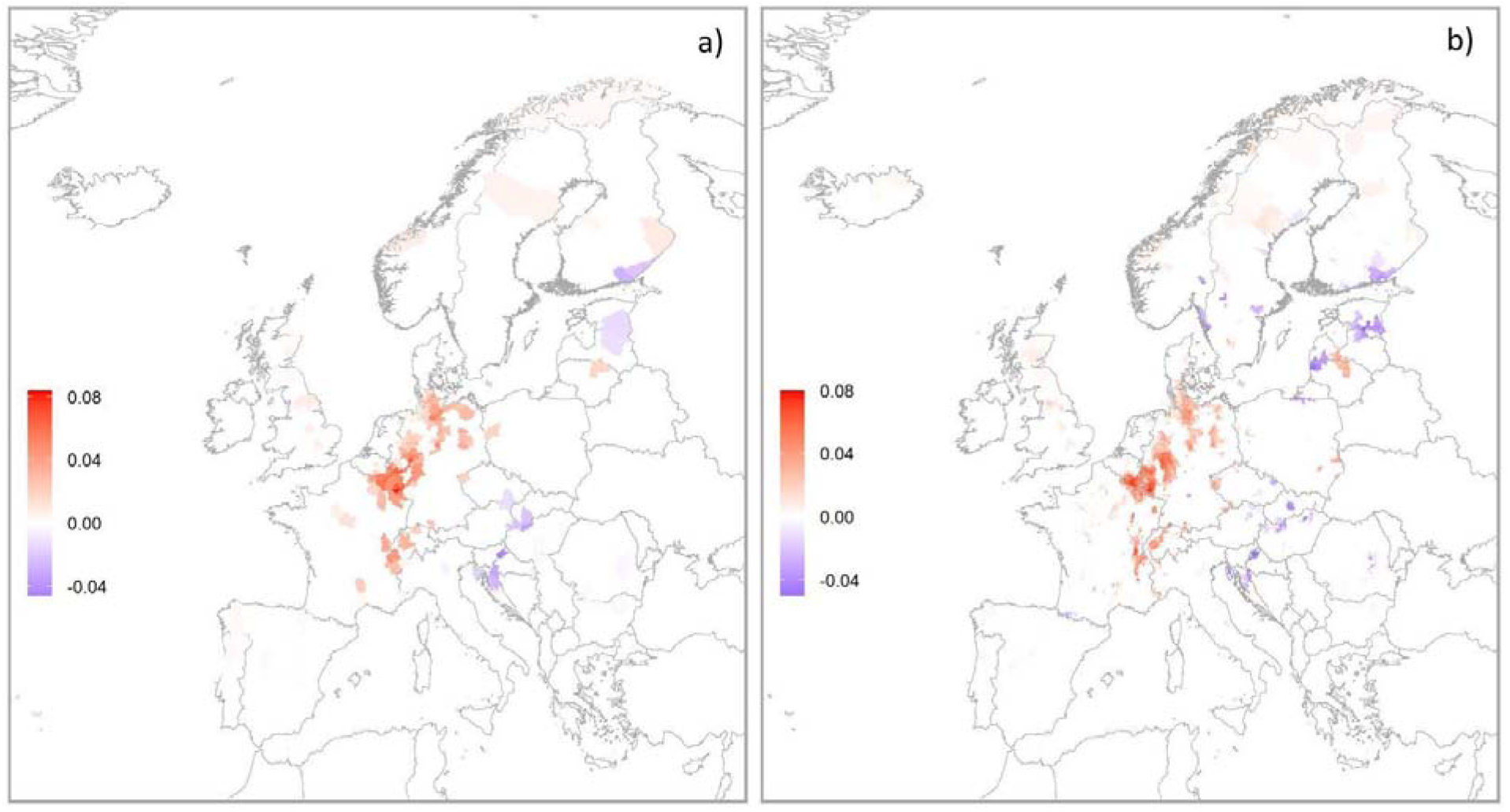
Significant coefficients (p < 0.05) of linear regression models applied to each area over 2017-2024 with year as predictor, at the NUTS-3 (a) and municipal (b) administrative levels. Only areas with statistically significant coefficients are shown in the maps.

## Discussion

The current spread and emergence of human TBE infections across Europe highlights the need to provide public health authorities with high-resolution predictive models as a tool to improve surveillance and preventive measures aimed at reducing the burden of TBE among European citizens. Here, we present a machine learning modelling framework that enables the assessment of the probability of changes in human TBE occurrence, on a yearly basis.

TBEV circulates in small, localized hotspots among ticks and hosts. However, due to the lack of centralized and consistent monitoring, comprehensive data on TBEV prevalence in wildlife and vectors are not available continent-wide. Therefore, in this study, we specifically trained our model on confirmed human TBE cases reported to ECDC at the NUTS-3 level from countries with mandatory reporting requirements. This choice provides a standardized, cross-border dataset that, while coarser in resolution, offers a robust basis for regional predictions across Europe. This decision underscores our focus on human TBE cases, rather than TBEV detections in wildlife or vectors, addressing the public health impact of TBE in human populations. For the same reason, we considered not only factors related to the natural hazard of TBE virus circulation, such as climate, vegetation, and presence of vertebrate hosts, which influence tick abundance and TBEV prevalence in ticks but also drivers related to human exposure to tick bites.

Overall, the spatio-temporal risk maps suggest a shift of human TBE suitability towards northern, western, and possibly also southwestern countries, with a significant positive trend in areas where TBEV is already recognised as endemic, such as Germany and Switzerland, and also in countries where so far only a few autochthonous cases have been detected, such as Belgium [36], France [37], and the UK [38]. Unfortunately, data from emerging TBE Countries, such as the UK, Belgium, and the Netherlands, were either unavailable in the ECDC TESSy dataset or not recorded at the NUTS-3 level, and thus were not included in the present study. To improve predictive accuracy for these areas, future studies should incorporate data from emerging regions as it becomes available. In any case, our results are consistent with the recent longitudinal and latitudinal shifts, which imply retraction from some areas as well as expansion into others, that were observed in cases reported to ECDC [6,40] but also suggest a possible new pathway of TBE emergence towards southwestern areas. We speculate these patterns could be driven by changes in climatic and ecological factors, such as the spatio-temporal expansion of ticks and their range of activity, coupled with host movements, changes in land use patterns and rise in exposure driven by changes in human behaviour and increase use of green areas also in relation to a warmer climate [41] (see Supplementary Material for a detailed explanation on covariates used in the model).

Our results highlight the relevance of including information on the occurrence of competent and tick-feeding hosts when modelling TBE. In small-scale studies, this is usually achieved by relying on locally recorded game population densities [19,42,43]. However, high-resolution host data are not available at the continental scale, which is why this information is generally not considered in European-wide studies. For this reason, we included in our modelling framework a variable describing the habitat suitability of critical host species, which was originally obtained using random forest and boosted regression trees-based spatial modelling techniques [25] and has already been shown to be a good predictor of TBE incidence across Europe [24]. A limitation of this study is that the variable used as a proxy for TBE host distribution does not consider the actual annual and interannual variations in host population density and is constant throughout the study period. However, it was found to significantly improve model performance compared to the same model built excluding this predictor (see Supplementary Material, Table S4, for the predictive performance of the BRT model without this variable).

Layers representing the distribution of the vectors, *Ixodes ricinus* and *Ixodes persulcatus* were initially considered as predictors but had no significant impact on the model’s performance and were therefore excluded from further analyses. This can be explained by the fact that the mere presence of the vector, which is widely distributed, is not sufficient for the circulation of the virus. Instead, specific ecological and environmental conditions are necessary.

Moreover, the incorporation of data on food-borne TBE infections, which were not included in this study due to the unavailability of information on the specific mode of TBEV transmission (either tick bites or infected food sources, such as unpasteurised milk and derived products), and data on the occurrence of cattle, sheep and goats, which act as sentinel and source of TBEV could enhance one-health surveillance. Yet, at this stage, our approach does not differentiate between tick-borne and food-borne human TBE infections, and the model indicates potential areas for disease transmission regardless of the infection mode. In the future, vaccination rates should also be considered to model changes in the true incidence of TBE in the human population, which we could not assess in this study due to the lack of available vaccination data coverage in all countries considered in this study.

We downscaled our results to the municipal administrative level to explore local patterns. These higher-resolution maps provide a more detailed picture of the pattern of human TBE suitable areas across Europe and could allow more effective and targeted surveillance efforts. The enhanced resolution of the municipal predictions shown in Figure 4 highlights the geographical heterogeneity of TBE occurrence over the years studied, which is consistent with the known patchy distribution of TBE infections across Europe. However, we were only able to validate these results using a limited dataset of recorded human TBE cases provided by five European countries (Austria, Finland, Italy, Lithuania, and Slovakia) upon special agreement. To facilitate future efforts to model human TBE occurrence at high spatial resolution, it would be beneficial to collect epidemiological data in an integrated and standardised manner across Europe, not only at the regional level but also at the municipality level.

Estimating and predicting the probability of occurrence of human TBE cases across Europe is essential for planning public health interventions, including awareness campaigns, vaccination programmes, and complementary risk assessments that are fundamental to inform and educate local populations in endemic regions, but also travellers, to take appropriate prophylactic measures to avoid tick bites and tick-borne infections.

In this study, we provide a solid framework for the assessment of areas where human TBE infections are likely to occur, which will be provided on an annual basis prior to the start of the tick questing season. The model is based on standardised and biologically consistent covariates, and we established its validity as a tool to strengthen and inform disease surveillance, control and prevention strategies aimed at mitigating the public health impact of TBE.

## Bibliography

1. Gritsun TS, Lashkevich VA, Gould EA. Tick-borne encephalitis. Antiviral Res. 2003 Jan;57(1– 2):129–46.

2. Lindquist L, Vapalahti O. Tick-borne encephalitis. The Lancet. 2008 May;371(9627):1861–71.

3. Martello E, Gillingham EL, Phalkey R, Vardavas C, Nikitara K, Bakonyi T, et al. Systematic review on the non-vectorial transmission of Tick-borne encephalitis virus (TBEv). Ticks Tick-Borne Dis. 2022 Nov;13(6):102028.

4. Ruzek D, Avšič Županc T, Borde J, Chrdle A, Eyer L, Karganova G, et al. Tick-borne encephalitis in Europe and Russia: Review of pathogenesis, clinical features, therapy, and vaccines. Antiviral Res. 2019 Apr;164:23–51.

5. European Centre for Disease Prevention and Control. Tick-borne encephalitis. In: ECDC. Annual epidemiological report for 2022. Stockholm: ECDC. 2024.

6. Van Heuverswyn J, Hallmaier-Wacker LK, Beauté J, Gomes Dias J, Haussig JM, Busch K, et al. Spatiotemporal spread of tick-borne encephalitis in the EU/EEA, 2012 to 2020. Eurosurveillance [Internet]. 2023 Mar 16 [cited 2023 Mar 20];28(11). Available from: https://www.eurosurveillance.org/content/10.2807/1560-7917.ES.2023.28.11.2200543

7. Elith J, Leathwick JR, Hastie T. A working guide to boosted regression trees. J Anim Ecol. 2008 Jul;77(4):802–13.

8. Bhatt S, Gething PW, Brady OJ, Messina JP, Farlow AW, Moyes CL, et al. The global distribution and burden of dengue. Nature. 2013 Apr;496(7446):504–7.

9. Nsoesie EO, Kraemer MU, Golding N, Pigott DM, Brady OJ, Moyes CL, et al. Global distribution and environmental suitability for chikungunya virus, 1952 to 2015. Eurosurveillance [Internet]. 2016 May 19 [cited 2024 Mar 13];21(20). Available from: https://www.eurosurveillance.org/content/10.2807/1560-7917.ES.2016.21.20.30234

10. Erazo D, Grant L, Ghisbain G, Marini G, Colón-González FJ, Wint W, et al. Contribution of climate change to the spatial expansion of West Nile virus in Europe. Nat Commun. 2024 Feb 8;15(1):1196.

11. Farooq Z, Rocklöv J, Wallin J, Abiri N, Sewe MO, Sjödin H, et al. Artificial intelligence to predict West Nile virus outbreaks with eco-climatic drivers. Lancet Reg Health - Eur. 2022 Jun;17:100370.

12. Arsevska E, Hengl T, Singleton DA, Noble PJM, Caminade C, Eneanya OA, et al. Risk factors for tick attachment in companion animals in Great Britain: a spatiotemporal analysis covering 2014–2021. Parasit Vectors. 2024 Jan 22;17(1):29.

13. Porretta D, Mastrantonio V, Amendolia S, Gaiarsa S, Epis S, Genchi C, et al. Effects of global changes on the climatic niche of the tick Ixodes ricinus inferred by species distribution modelling. Parasit Vectors. 2013 Dec;6(1):271.

14. Walter M, Vogelgesang JR, Rubel F, Brugger K. Tick-Borne Encephalitis Virus and Its European Distribution in Ticks and Endothermic Mammals. Microorganisms. 2020 Jul 17;8(7):1065.

15. Borde JP, Glaser R, Braun K, Riach N, Hologa R, Kaier K, et al. Decoding the Geography of Natural TBEV Microfoci in Germany: A Geostatistical Approach Based on Land-Use Patterns and Climatological Conditions. Int J Environ Res Public Health. 2022 Sep 19;19(18):11830.

16. Uusitalo R, Siljander M, Dub T, Sane J, Sormunen JJ, Pellikka P, et al. Modelling habitat suitability for occurrence of human tick-borne encephalitis (TBE) cases in Finland. Ticks Tick-Borne Dis. 2020 Sep;11(5):101457.

17. Friedsam AM, Brady OJ, Pilic A, Dobler G, Hellenbrand W, Nygren TM. Geo-Spatial Characteristics of 567 Places of Tick-Borne Encephalitis Infection in Southern Germany, 2018– 2020. Microorganisms. 2022 Mar 17;10(3):643.

18. Zeimes CB, Olsson GE, Hjertqvist M, Vanwambeke SO. Shaping zoonosis risk: landscape ecology vs. landscape attractiveness for people, the case of tick-borne encephalitis in Sweden. Parasit Vectors. 2014;7(1):370.

19. Rizzoli A, Hauffe HC, Tagliapietra V, Neteler M, Rosà R. Forest Structure and Roe Deer Abundance Predict Tick-Borne Encephalitis Risk in Italy. Moen J, editor. PLoS ONE. 2009 Feb 2;4(2):e4336.

20. Tkadlec E, Václavík T, Široký P. Rodent Host Abundance and Climate Variability as Predictors of Tickborne Disease Risk 1 Year in Advance. Emerg Infect Dis. 2019 Sep;25(9):1738–41.

21. Saegerman C, Humblet MF, Leandri M, Gonzalez G, Heyman P, Sprong H, et al. First Expert Elicitation of Knowledge on Possible Drivers of Observed Increasing Human Cases of Tick-Borne Encephalitis in Europe. Viruses. 2023 Mar 20;15(3):791.

22. Regulation (EC) No 1059/2003 of the European Parliament and of the Council of 26 May 2003 on the establishment of a common classification of territorial units for statistics (NUTS) [Internet]. OJ L May 26, 2003. Available from: http://data.europa.eu/eli/reg/2003/1059/oj/eng

23. Braks MAH, Van Wieren SE, Takken W, Sprong H, editors. Ecology and prevention of Lyme borreliosis. Wageningen: Wageningen Academic Publishers; 2016. 462 p. (Ecology and control of vector-borne diseases / ed. by: Willem Takken).

24. Dagostin F, Tagliapietra V, Marini G, Cataldo C, Bellenghi M, Pizzarelli S, et al. Ecological and environmental factors affecting the risk of tick-borne encephalitis in Europe, 2017 to 2021. Eurosurveillance [Internet]. 2023 Oct 19 [cited 2023 Nov 7];28(42). Available from: https://www.eurosurveillance.org/content/10.2807/1560-7917.ES.2023.28.42.2300121

25. Wint W, Morley D, Jolyon Medlock, Alexander N. A first attempt at modelling red deer (Cervus elaphus) distributions over Europe [Internet]. figshare; 2014 [cited 2022 Apr 11]. Available from: https://figshare.com/articles/dataset/A_first_attempt_at_modelling_red_deer_Cervus_elaphus_distributions_over_Europe/1008334/1

26. Tatem AJ. WorldPop, open data for spatial demography. https://www.worldpop.org. Sci Data. 2017 Dec;4(1):170004.

27. G. Ridgeway, Edwards D, Kriegler B, Schroedl S, Soutworth H, Greenwell B, et al. gbm: Generalized Boosted Regression Models. R package version 2.1.3. [Internet]. 2017. Available from: Available from: https://cran.r-project.org/package=gbm.

28. Lang M, Binder M, Richter J, Schratz P, Pfisterer F, Coors S, et al. mlr3: A modern object-oriented machine learning framework in R. J Open Source Softw. 2019 Dec 11;4(44):1903.

29. Valavi R, Elith J, Lahoz-Monfort JJ, Guillera-Arroita G. BLOCK CV ⍰: An R package for generating spatially or environmentally separated folds for k -fold cross-validation of species distribution models. Warton D, editor. Methods Ecol Evol. 2019 Feb;10(2):225–32.

30. Leroy B, Delsol R, Hugueny B, Meynard CN, Barhoumi C, Barbet-Massin M, et al. Without quality presence–absence data, discrimination metrics such as TSS can be misleading measures of model performance. J Biogeogr. 2018 Sep;45(9):1994–2002.

31. Sørensen T. A Method of Establishing Groups of Equal Amplitude in Plant Sociology Based on Similarity of Species and Its Application to Analyses of the Vegetation on Danish Commons. K Dan Vidensk Selsk. 1948;(5):1–34.

32. Jiménez-Valverde A. Insights into the area under the receiver operating characteristic curve (AUC) as a discrimination measure in species distribution modelling. Glob Ecol Biogeogr. 2012 Apr;21(4):498–507.

33. Ghisbain G, Thiery W, Massonnet F, Erazo D, Rasmont P, Michez D, et al. Projected decline in European bumblebee populations in the twenty-first century. Nature. 2024 Apr 11;628(8007):337–41.

34. Da Re D, Gilbert M, Chaiban C, Bourguignon P, Thanapongtharm W, Robinson TP, et al. Downscaling livestock census data using multivariate predictive models: Sensitivity to modifiable areal unit problem. Koukoulas S, editor. PLOS ONE. 2020 Jan 27;15(1):e0221070.

35. R Core Team. R: A language and environment for statistical computing. R Foundation for Statistical Computing. [Internet]. Vienna, Austria; 2022. Available from: https://www.R-project.org/

36. Stoefs A, Heyndrickx L, De Winter J, Coeckelbergh E, Willekens B, Alonso-Jiménez A, et al. Autochthonous Cases of Tick-Borne Encephalitis, Belgium, 2020. Emerg Infect Dis. 2021 Aug;27(8):2179–82.

37. Botelho-Nevers E, Gagneux-Brunon A, Velay A, Guerbois-Galla M, Grard G, Bretagne C, et al. Tick-Borne Encephalitis in Auvergne-Rhône-Alpes Region, France, 2017–2018. Emerg Infect Dis. 2019 Oct;25(10):1944–8.

38. Kreusch TM, Holding M, Hewson R, Harder T, Medlock JM, Hansford KM, et al. A probable case of tick-borne encephalitis (TBE) acquired in England, July 2019. Euro Surveill Bull Eur Sur Mal Transm Eur Commun Dis Bull. 2019 Nov;24(47).

39. UK Health Security Agency. Tick-borne encephalitis detection in England [Press release]. 2023 May 4; Available from: https://www.gov.uk/government/news/tick-borne-encephalitis-detection-in-england

40. European Centre for Disease Prevention and Control. Tick-borne encephalitis. In: ECDC. Annual epidemiological report for 2020. Stockholm: ECDC. 2022.

41. Bille RA, Jensen KE, Buitenwerf R. Global patterns in urban green space are strongly linked to human development and population density. Urban For Urban Green. 2023 Aug;86:127980.

42. Dub T, Ollgren J, Huusko S, Uusitalo R, Siljander M, Vapalahti O, et al. Game Animal Density, Climate, and Tick-Borne Encephalitis in Finland, 2007–2017. Emerg Infect Dis. 2020 Dec;26(12):2899–906.

43. Kriz B, Daniel M, Benes C, Maly M. The Role of Game (Wild Boar and Roe Deer) in the Spread of Tick-Borne Encephalitis in the Czech Republic. Vector-Borne Zoonotic Dis. 2014 Nov;14(11):801–7.

44. Achazi K, RůŽek D, Donoso-Mantke O, Schlegel M, Ali HS, Wenk M, et al. Rodents as Sentinels for the Prevalence of Tick-Borne Encephalitis Virus. Vector-Borne Zoonotic Dis. 2011 Jun;11(6):641–7.

45. Bolzoni L, Rosà R, Cagnacci F, Rizzoli A. Effect of deer density on tick infestation of rodents and the hazard of tick-borne encephalitis. II: Population and infection models. Int J Parasitol. 2012 Apr;42(4):373–81.

46. Cagnacci F, Bolzoni L, Rosà R, Carpi G, Hauffe HC, Valent M, et al. Effects of deer density on tick infestation of rodents and the hazard of tick-borne encephalitis. I: Empirical assessment. Int J Parasitol. 2012 Apr;42(4):365–72.

47. Knap N, Avšič-Županc T. Correlation of TBE Incidence with Red Deer and Roe Deer Abundance in Slovenia. Yu X jie, editor. PLoS ONE. 2013 Jun 11;8(6):e66380.

48. Dagostin F, Tagliapietra V, Marini G, Ferrari G, Cervellini M, Wint W, et al. High habitat richness reduces the risk of tick-borne encephalitis in Europe: A multi-scale study. One Health. 2024 Jun;18:100669.

49. Ostfeld RS, Holt RD. Are predators good for your health? Evaluating evidence for top-down regulation of zoonotic disease reservoirs. Front Ecol Environ. 2004 Feb;2(1):13–20.

50. Rosà R. Changes in host densities and co-feeding pattern efficiently predict tick-borne encephalitis hazard in an endemic focus in northern Italy. Int J Parasitol. 2019.

