## Supplementary materials for "Modeling spatio-temporal annual changes in the probability of human tick-borne encephalitis (TBE) occurrence across Europe"

SUPPLEMENTARY MATERIAL

**Description of covariates and data sources**

To model tick-borne encephalitis (TBE) risk of human infections across Europe, we selected key hazard factors suggested by Dagostin and colleagues [1]. For each spatial unit, we extracted the percentage of forested areas (Figure S1) from the 2018 Corine Land Cover (CLC) data inventory (class “3.1”) with a resolution of 0.25x0.25 km [2]. We used satellite images acquired by the Moderate Resolution Imaging Spectroradiometer (MODIS) and supplied by NASA with a resolution of 5.6 km as a source of Land Surface Temperature (LST) [3], and the ECMWF ERA5-Land dataset at 30 arc seconds resolution as a source of cumulative precipitation data [4]. We calculated the total precipitation of the driest quarter of the previous year (Figure S2) following the formula stated in the World Climate (WorldClim) Database [5] and computed the autumnal cooling rate of the previous year (Figure S3) by applying a linear regression to the average daily temperature against the Julian day in the period 1st August – 31st October [6]. We computed the density of forest roads contained by OpenStreetMap as a proxy of accessibility to forested areas (Figure S4). Human population density in each area was derived from the WorldPop dataset (Figure S5). All explanatory variables were computed by averaging them for each administrative area included in the dataset.

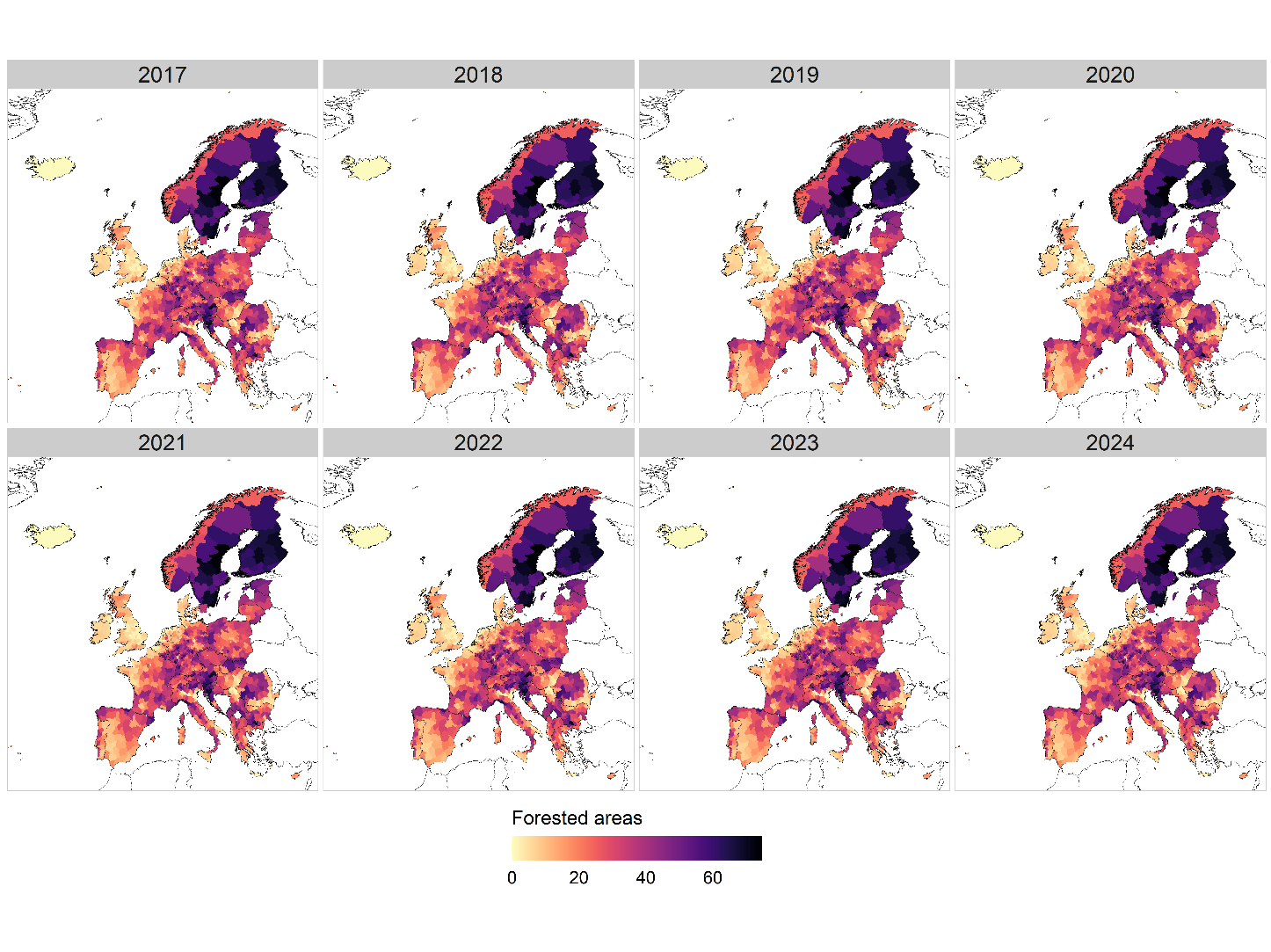

Figure S1 – Percentage of forested areas for each NUTS-3 administrative area (2017-2024).

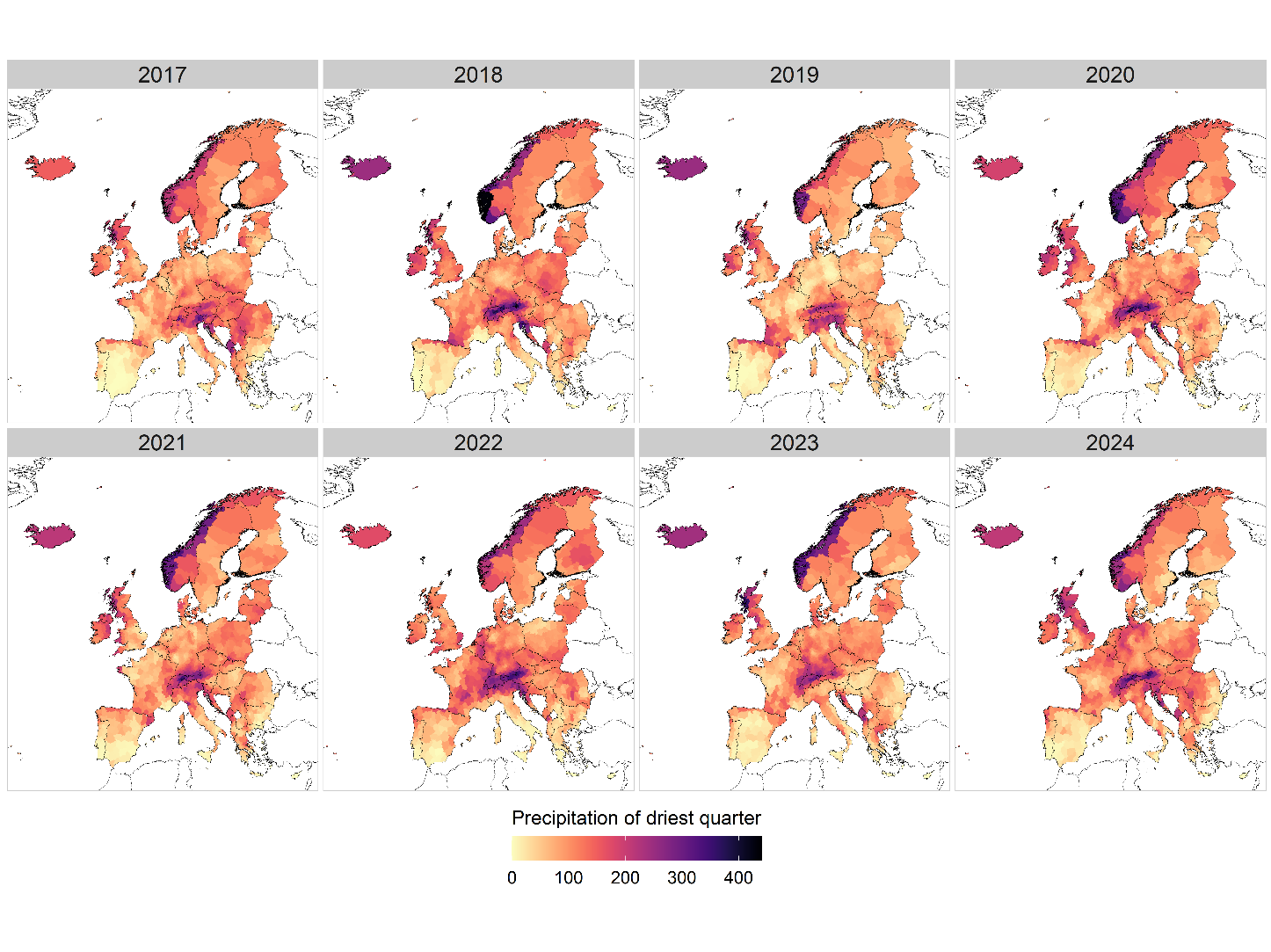

Figure S2 – Precipitation recorded in the driest quarter of the year for each NUTS-3 administrative area (2017-2024).

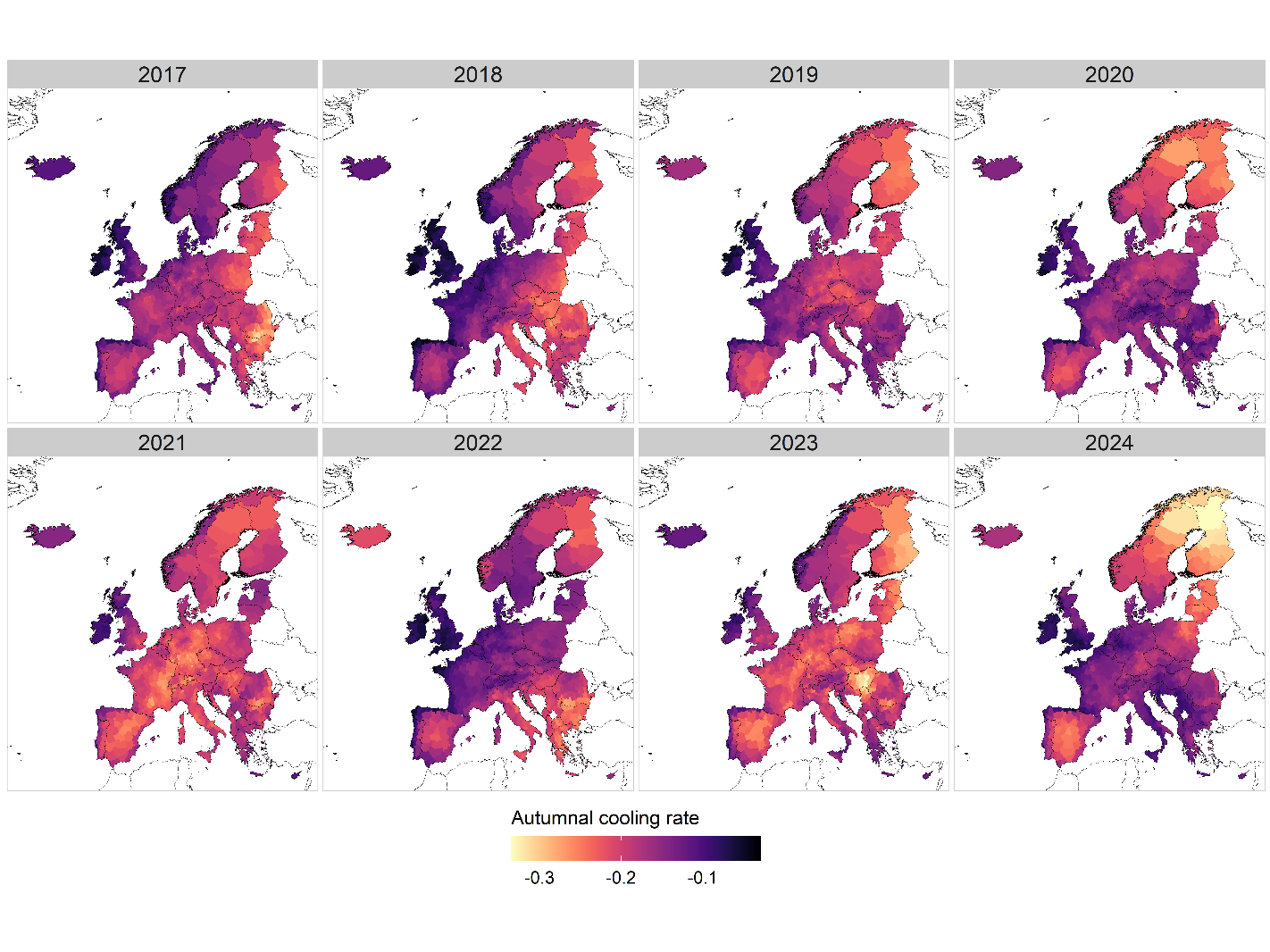

Figure S3 – Autumnal cooling rate for each NUTS-3 administrative area (2017-2024).

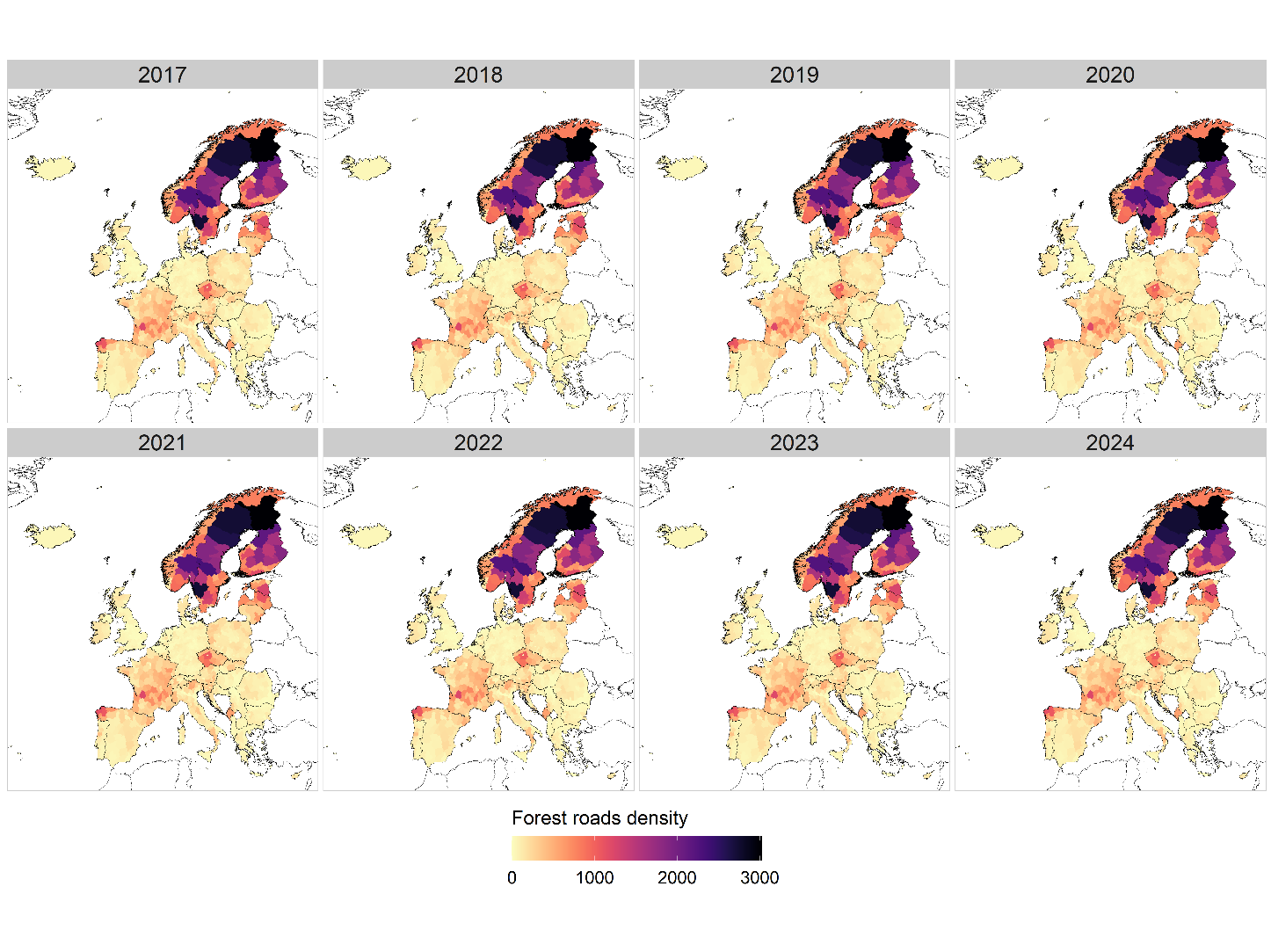

Figure S4 – Forest roads density for each NUTS-3 administrative area (2017-2024).

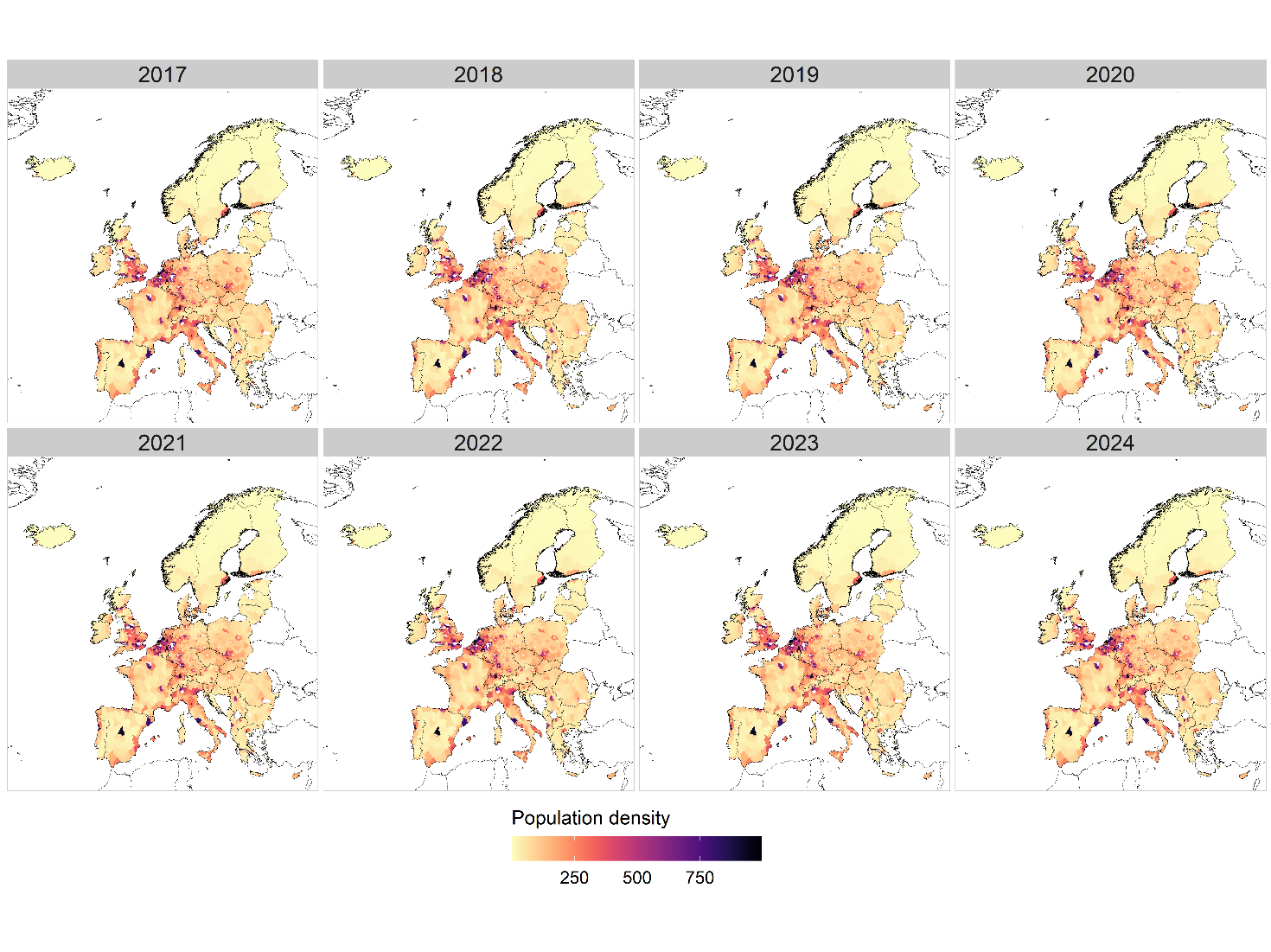
Figure S5 – Population density in each NUTS-3 administrative area (2017-2024).

To account for TBE vector hosts distribution, we used 1-km data about the weighted probability of presence of selected critical reservoir species (*Apodemus flavicollis*, *Myodes glareolus*), vector amplifiers species (*Dama dama*, *Cervus elaphus*, *Capreolus capreolus*, *Alces alces*, *Odocoileus virginianus*, *Lepus europaeus*, *Lepus timidus*), and predators (*Vulpes vulpes*) [7]. Specifically, the distributions of the known TBE vector hosts were obtained from IUCN and the European Mammal Atlas and converted to presence or absence within standard 50 km grid and then ranked from 1 to 4 according to their function as follows: (i) Reservoir Host and virus amplifiers (*Apodemus flavicollis*, *Myodes glareolus*), Score 4; (ii) Major vector amplifiers and virus dilution (*Capreolus capreolus*, *cervus elephus, Dama dama),* Score 3; (iii) Minor vector amplifiers and virus dilution (*Alces alces*, *Odocoelus virginianus*) and possible vector amplifiers (*Lepus europeus*, *Lepus timidus*), Score 2; Host predators (*vulpes vulpes*), Score -1. An equal number of zeros (“hosts absence”) points were assigned randomly outside the known host ranges. The weighted host presence was used as response variable and modelled using random forest and boosted regression trees approaches, implemented using the VECMAP modelling suite, using a standard set of covariates including Fourier Processed Remotely Sensed environmental variables. The outputs of the two methods were then ensembled as a simple mean (Figure S6). Table S1 and S2 show the top predictors for each method.

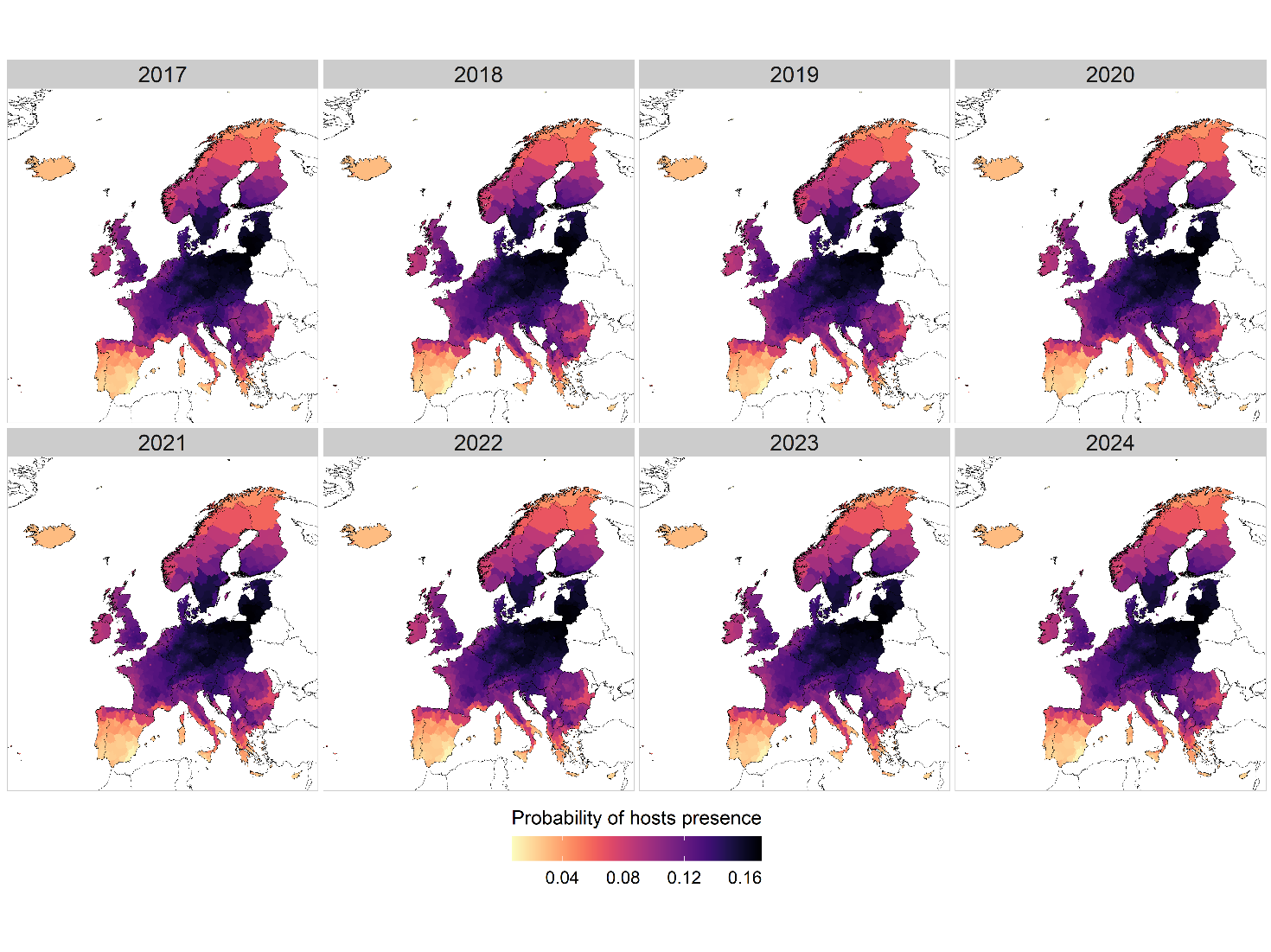

Figure S6 – Probability of hosts presence for each NUTS-3 administrative area (2017-2024).

| **Variable** | **%IncMSE** |
| --- | --- |
| Minimum rainfall | 10·55065 |
| EVI maximum | 4·11796 |
| Daytime LST maximum | 5·154958 |
| Mean rainfall | 4·439648 |
| NDVI maximum | 3·761641 |
| NDVI mean | 2·827918 |
| EVI mean | 3·110508 |
| Maximum rainfall | 1·780188 |
| Daytime LST mean | 2·229086 |
| Daytime LST phase 3  timing of peak of component 3 | 1·088382 |

Table S1 – Top predictors for the random forest model used to derive hosts suitability.

| **Variable** | **%IncMSE** |
| --- | --- |
| Minimum rainfall | 49·91 |
| Daytime LST maximum | 4·86 |
| NDVI mean | 4·07 |
| Daytime LST mean | 3·90 |
| Nighttime LST mean | 3·39 |
| Phase 1 timing of fourier annual component peak | 2·67 |
| NDVI maximum | 2·56 |
| Nighttime LST % var. annual cycle | 2·03 |
| GRUMP human Population density | 1·60 |
| Nighttime LST minimum | 1·48 |

 Table S2 – Top predictors for the boosted regression tree model used to derive hosts suitability.

**Presence and absence of human TBE cases at the municipal administrative level (2017-2022)**

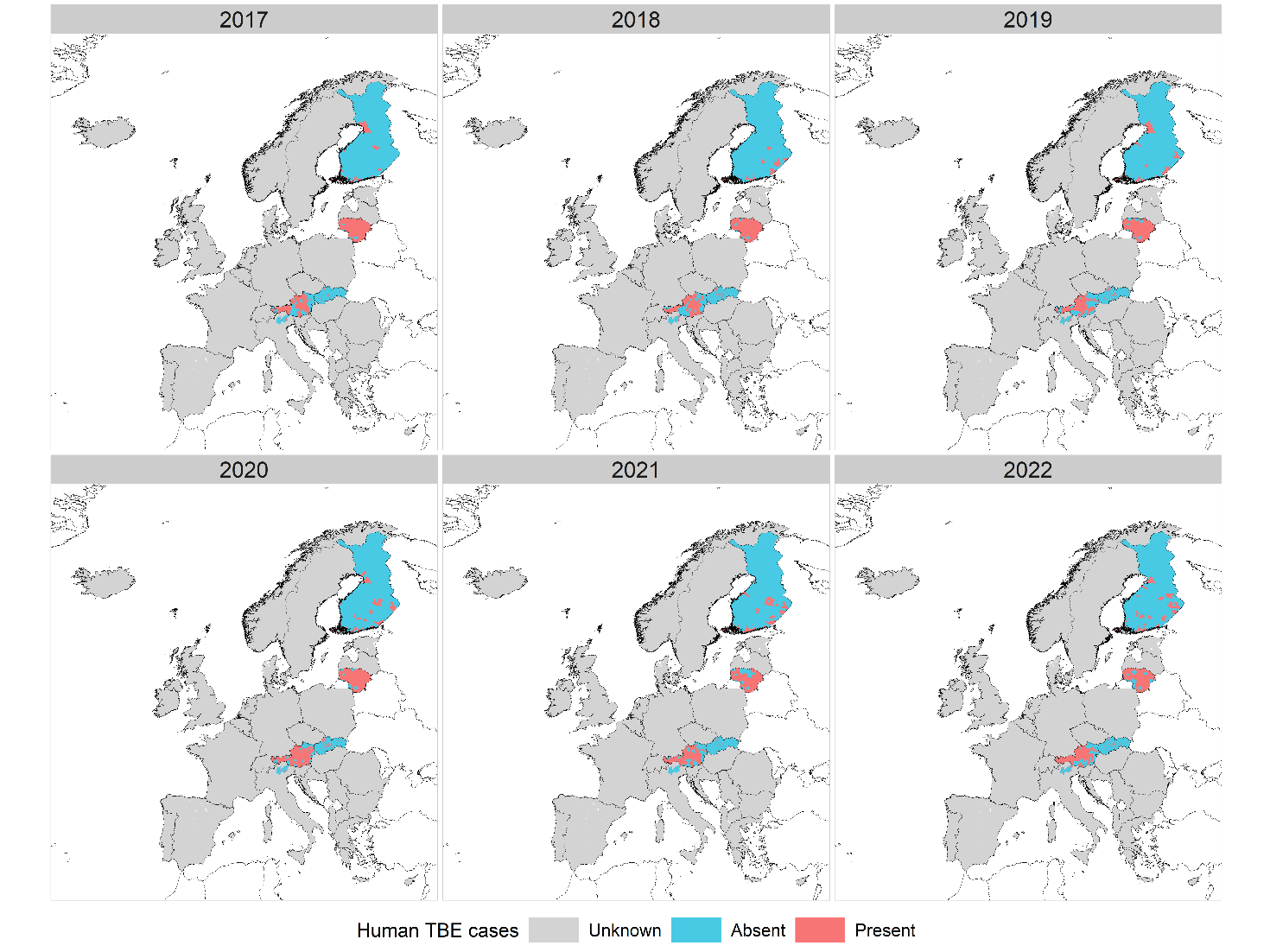

Figure S7 – Presence (in red) and absence (in blue) of human TBE cases at the municipal administrative level (2017-2022) derived from data provided upon agreement by Austria, Finland, Lithuania, Slovakia, Slovenia, and the Italian provinces of Trento and Belluno. Areas where TBE distribution is unknown at the municipal level are masked in grey.

**Predictive performance of the boosted regression trees (BRT) models**

| **Replicate** | **Area Under the Curve (AUC)** | **Sensitivity** | **Specificity** | **SI_ppc_ (threshold)** |
| --- | --- | --- | --- | --- |
| 1 | 0·82 | 0·83 | 0·80 | 0·70 (0·34) |
| 2 | 0·87 | 0·84 | 0·80 | 0·71 (0·32) |
| 3 | 0·86 | 0·81 | 0·82 | 0·71 (0·36) |
| 4 | 0·87 | 0·83 | 0·80 | 0·71 (0·34) |
| 5 | 0·82 | 0·82 | 0·81 | 0·71 (0·35) |
| 6 | 0·86 | 0·83 | 0·80 | 0·70 (0·31) |
| 7 | 0·86 | 0·82 | 0·81 | 0·71 (0·32) |
| 8 | 0·85 | 0·82 | 0·82 | 0·71 (0·33) |
| 9 | 0·85 | 0·82 | 0·82 | 0·71 (0·32) |
| 10 | 0·85 | 0·83 | 0·80 | 0·71 (0·34) |

Table S3 - Area under the curve (AUC), sensitivity, specificity, and prevalence-pseudoabsence-calibrated Sørensen index (SI_ppc_) values computed for each of the ten independent replicate boosted regression tree (BRT) models trained on the observed distribution of TBE human cases at NUTS-3 administrative level along the period 2017-2021. The SI_ppc_ were computed while performing an optimisation of the probability threshold in the range [0-1] with a 0.01 step increment. The threshold value maximising the SI_ppc_ is reported under parentheses.

**Partial dependence plots**

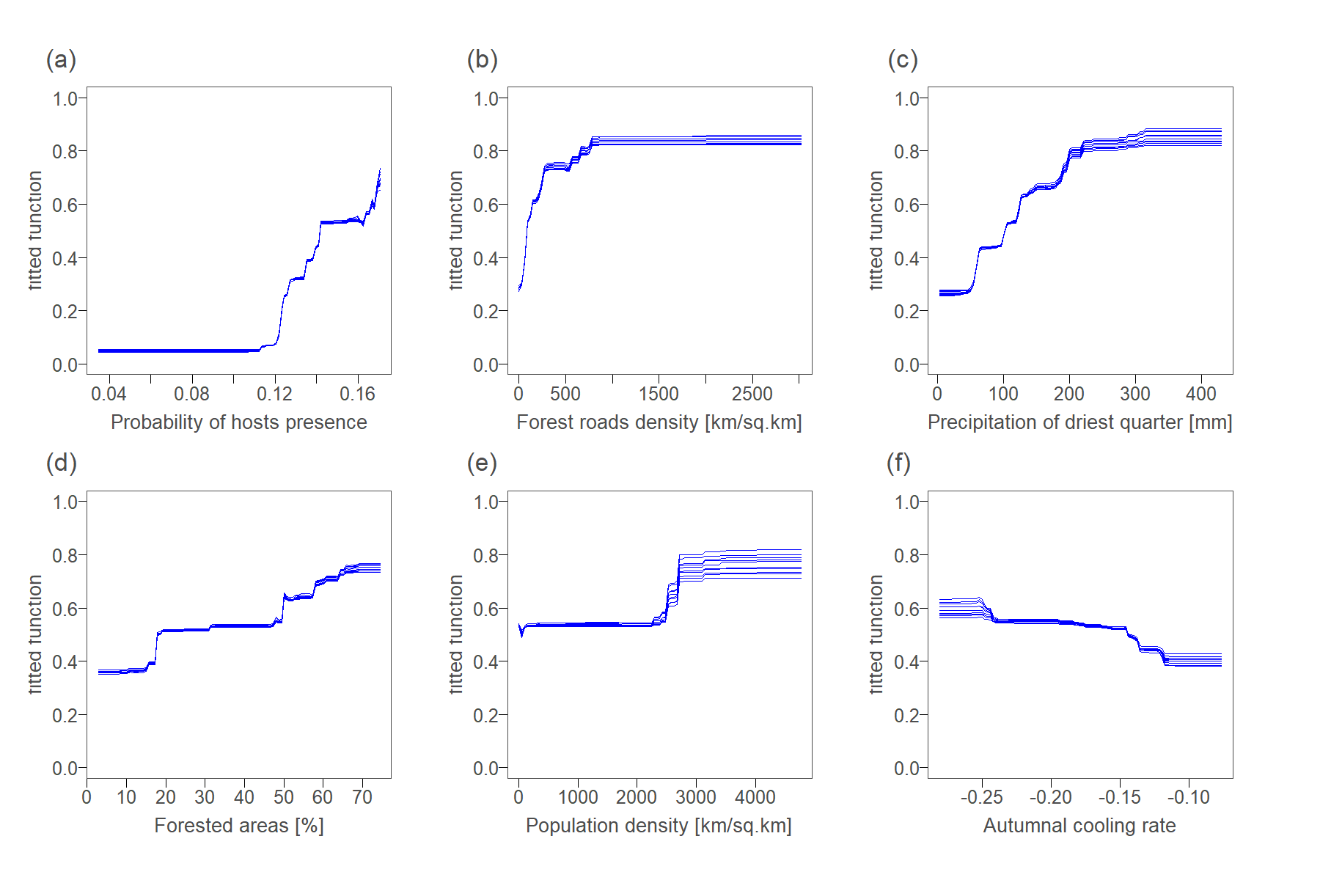

Figure S8. Fitted functions of each variable across the 10 replicates of the BRT model: (a) Probability of hosts presence (RI = 39.44); (b) Forest roads density (RI = 21.39); (c) Precipitation of the driest quarter (RI = 20.19); (d) Percentage of forested areas (RI = 14.85); (e) Population density (RI = 2.37); (f) Autumnal cooling rate (RI = 1.77).

**Additional discussion on the influence of variables on model predictions**

Abiotic factors directly affect the off-host stages of ticks, particularly temperature and humidity, which influence tick body hydration [8]. Our results suggest that areas with higher precipitation during the driest quarter of the year lead to higher TBE suitability, as rainfall has an indirect positive effect on relative humidity (and therefore on saturation deficit) that enhances tick survival and favours questing also during the driest months of the year [9]. Moreover, a high rate of autumnal cooling (i.e. a rapid drop in late summer temperatures) affects the critical seasonal point of diapause induction in ticks by favouring a behavioural diapause in larvae and nymphs that are ready for synchronised questing the following spring. This scenario enhances TBEV transmission from infected nymphs to co-feeding uninfected ticks on rodents, supporting TBEV circulation [10–12]. Concerning environmental variables, forests provide suitable habitats and resources for tick hosts, facilitating their encounter and resulting in higher TBE suitability [13,14]. Finally, human activity and behaviour can act in synergy with these ecological factors by increasing the likelihood of encountering infected ticks, as people who engage in recreational or occupational outdoor activities in green areas are at increased risk of tick bites [15,16]. Therefore, forest road density and population density, included in the model as proxies for forest accessibility and human exposure to infected ticks, are both associated with higher probabilities of human infections with TBE. Our results also highlight the importance of considering the presence of key hosts when modelling TBE. Rodent species, such as *Apodemus flavicollis* and *Myodes glareolus*, directly support viral circulation by transmitting the virus to feeding ticks. This ongoing transmission between rodents and ticks helps maintain a steady presence of the virus within tick populations [12]. In addition, deer species (*Cervus elaphus*, *Dama dama*, and *Capreolus capreolus*) amplify tick abundance by providing feeding opportunities. Deer also contribute to the spread of ticks over wider areas as they migrate or move across their ranges, facilitating tick colonization in new locations [13].

**Predictive performance of the boosted regression trees (BRT) models excluding the predictor “probability of hosts presence”**

| **Replicate** | **Area Under the Curve (AUC)** | **Sensitivity** | **Specificity** | **SI_ppc_ (threshold)** |
| --- | --- | --- | --- | --- |
| 1 | 0·76 | 0·79 | 0·72 | 0·64 (0.30) |
| 2 | 0·76 | 0·78 | 0·72 | 0·63 (0.30) |
| 3 | 0·79 | 0·79 | 0·72 | 0·64 (0.30) |
| 4 | 0·77 | 0·78 | 0·72 | 0·63 (0.30) |
| 5 | 0·78 | 0·78 | 0·72 | 0·63 (0.30) |
| 6 | 0·76 | 0·79 | 0·73 | 0·64 (0.30) |
| 7 | 0·76 | 0·77 | 0·73 | 0·63 (0.30) |
| 8 | 0·76 | 0·79 | 0·72 | 0·64 (0.30) |
| 9 | 0·79 | 0·79 | 0·72 | 0·63 (0.30) |
| 10 | 0·78 | 0·79 | 0·72 | 0·63 (0.30) |

Table S4 - Area under the curve (AUC), sensitivity, specificity, and prevalence-pseudoabsence-calibrated Sørensen index (SI_ppc_) values computed for each of the ten independent replicate boosted regression tree (BRT) models trained on the observed distribution of TBE human cases at NUTS-3 administrative level along the period 2017-2021, excluding the “probability of hosts presence” predictor. The S_Ippc_ were computed while performing an optimisation of the probability threshold in the range [0-1] with a 0.01 step increment. The threshold value maximising the SI_ppc_ is reported under parentheses.

**Bibliography**

1.	Dagostin F, Tagliapietra V, Marini G, Cataldo C, Bellenghi M, Pizzarelli S, et al. Ecological and environmental factors affecting the risk of tick-borne encephalitis in Europe, 2017 to 2021. Eurosurveillance [Internet]. 2023 Oct 19 [cited 2023 Nov 7];28(42). Available from: https://www.eurosurveillance.org/content/10.2807/1560-7917.ES.2023.28.42.2300121

2.	Feranec J. Project CORINE land cover. In: European Landscape Dynamics: CORINE Land Cover Data. 2016. p. 9–14.

3.	Wan Z, Hook S, Hulley G. MODIS/Terra Land Surface Temperature/Emissivity 8-Day L3 Global 0.05Deg CMG V061 [Internet]. NASA EOSDIS Land Processes DAAC; 2021 [cited 2022 Sep 1]. Available from: https://lpdaac.usgs.gov/products/mod11c2v061/

4.	Copernicus Climate Change Service. ERA5-Land hourly data from 2001 to present [Internet]. ECMWF; 2019 [cited 2022 Apr 11]. Available from: https://cds.climate.copernicus.eu/doi/10.24381/cds.e2161bac

5.	O’Donnell MS, Ignizio DA. Bioclimatic predictors for supporting ecological applications in the conterminous United States. U.S. Geological Survey Data Series, 691 10 p. 2012. 691 10 p.

6.	Randolph SE, Green RM, Peacey MF, Rogers DJ. Seasonal synchrony: the key to tick-borne encephalitis foci identified by satellite data. Parasitology. 2000 Jul;121(1):15–23.

7.	Wint W, Morley D, Jolyon Medlock, Alexander N. A first attempt at modelling red deer (Cervus elaphus) distributions over Europe [Internet]. figshare; 2014 [cited 2022 Apr 11]. Available from: https://figshare.com/articles/dataset/A_first_attempt_at_modelling_red_deer_Cervus_elaphus_distributions_over_Europe/1008334/1

8.	Borde JP, Kaier K, Hehn P, Matzarakis A, Frey S, Bestehorn M, et al. The complex interplay of climate, TBEV vector dynamics and TBEV infection rates in ticks—Monitoring a natural TBEV focus in Germany, 2009–2018. Forrester N, editor. PLOS ONE. 2021 Jan 7;16(1):e0244668.

9.	Porretta D, Mastrantonio V, Amendolia S, Gaiarsa S, Epis S, Genchi C, et al. Effects of global changes on the climatic niche of the tick Ixodes ricinus inferred by species distribution modelling. Parasit Vectors. 2013 Dec;6(1):271.

10.	Carpi G, Cagnacci F, Neteler M, Rizzoli A. Tick infestation on roe deer in relation to geographic and remotely sensed climatic variables in a tick-borne encephalitis endemic area. Epidemiol Infect. 2008 Oct;136(10):1416–24.

11.	Rosà R, Andreo V, Tagliapietra V, Baráková I, Arnoldi D, Hauffe H, et al. Effect of Climate and Land Use on the Spatio-Temporal Variability of Tick-Borne Bacteria in Europe. Int J Environ Res Public Health. 2018 Apr 12;15(4):732.

12.	Rosà R, Tagliapietra V, Manica M, Arnoldi D, Hauffe HC, Rossi C, et al. Changes in host densities and co-feeding pattern efficiently predict tick-borne encephalitis hazard in an endemic focus in northern Italy. Int J Parasitol. 2019 Sep;49(10):779–87.

13.	Rizzoli A, Hauffe HC, Tagliapietra V, Neteler M, Rosà R. Forest Structure and Roe Deer Abundance Predict Tick-Borne Encephalitis Risk in Italy. Moen J, editor. PLoS ONE. 2009 Feb 2;4(2):e4336.

14.	Vanwambeke SO, Sumilo D, Bormane A, Lambin EF, Randolph SE. Landscape predictors of tick-borne encephalitis in Latvia: land cover, land use, and land ownership. Vector Borne Zoonotic Dis Larchmt N. 2010 Jun;10(5):497–506.

15.	Stefanoff P, Rosinska M, Samuels S, White DJ, Morse DL, Randolph SE. A National Case-Control Study Identifies Human Socio-Economic Status and Activities as Risk Factors for Tick-Borne Encephalitis in Poland. Munderloh UG, editor. PLoS ONE. 2012 Sep 19;7(9):e45511.

16.	Zeimes CB, Olsson GE, Hjertqvist M, Vanwambeke SO. Shaping zoonosis risk: landscape ecology vs. landscape attractiveness for people, the case of tick-borne encephalitis in Sweden. Parasit Vectors. 2014;7(1):370.
